# CRISPRi-seq in *Haemophilus influenzae* reveals genome-wide and medium-specific growth determinants

**DOI:** 10.1101/2025.08.05.668841

**Authors:** Celia Gil-Campillo, Johann Mignolet, Asier Domínguez-San Pedro, Beatriz Rapún- Araiz, Axel B. Janssen, Vincent de Bakker, Jan-Willem Veening, Junkal Garmendia

**Author notes:** Corresponding authors: Junkal Garmendia, Instituto de Agrobiotecnología, CSIC-Gobierno de Navarra, 31192 Mutilva, Navarra, Spain;, Jan-Willem Veening, Department of Fundamental Microbiology, Faculty of Biology and Medicine, University of Lausanne, Biophore Building, 1015, Lausanne, Switzerland. Equal contribution (Celia Gil-Campillo, Johann Mignolet). **Declarations of interest:** J.W.V. is a scientific advisory board member at i-Seq Biotechnology.

## Abstract

Work in the human pathobiont *Haemophilus influenzae* has pioneered functional genomics in bacteria such as genome-wide transposon mutagenesis combined with deep sequencing. These approaches unveiled a large set of likely essential genes, but functional studies are hampered due to a limited molecular toolbox. To bridge this gap, we engineered a titratable anhydrotetracycline (aTc)-inducible CRISPRi (Clustered Regularly Interspaced Short Palindromic Repeats interference) platform for efficient regulation of gene expression in *H. influenzae*. Genome-wide fitness analyses in two different *in vitro* culture media by CRISPRi-seq revealed growth medium-dependent fitness cost for a panel of *H. influenzae* genes. We demonstrated that CRISPRi-programmed fitness defects can be rescuable, and refined previous Tn-seq based essentialome studies. Finally, we introduce HaemoBrowse, an extensive user-friendly online resource for visual inspection of *H. influenzae* genome annotations, including sgRNA spacers. The inducible CRISPRi platform described here represents a valuable tool enabling functional genomics and the study of essential genes, thereby contributing to the identification of therapeutic targets for developing drugs and vaccines against *H. influenzae*.

**Importance:** CRISPRi-seq is a robust method to study bacterial gene fitness and essentiality via relative quantification and comparison of sgRNA abundance at a genome-wide scale. Here, we present a novel CRISPRi system for individual genes or pooled libraries knockdown in *Haemophilus influenzae*. A genome-wide CRISPRi library designed to cover 99.27% of all total genetic features in the genome of RdKW20 strain was constructed and screened in two laboratory growth media through CRISPRi-seq, uncovering growth medium-dependent fitness cost, further confirmed with individual knockdown/knockout mutants. We also introduce HaemoBrowse (https://HaemoBrowse.VeeningLab.com), through which genome annotations and sgRNA design on *H. influenzae* genomes can be readily inspected. This platform provides a valuable tool for gene function and essentiality analyses in a notorious human pathobiont.

## Introduction

The human-specific Gram-negative bacterial species *Haemophilus influenzae* is a normal part of the upper airway microbiome but can cause invasive infections such as pneumonia, otitis media, and meningitis typically in infants and young children. Infant mortality due to invasive infections was drastically reduced by introduction of the conjugated vaccine that targets strains with the serotype *b* polysaccharide capsule (1). However, nontypeable *H. influenzae* strains continue to cause high morbidity due to their role in common infections and chronic diseases, and are major contributors to persistent infection and exacerbations of chronic obstructive pulmonary disease (COPD) and cystic fibrosis (CF) (2–4).

Functional genomics is a powerful approach to identify new vaccine candidates and antibiotic targets for important human pathogens. Genome-wide studies to assess the fitness contribution or essentiality of each genetic feature across diverse conditions or genetic backgrounds in bacteria have predominantly been performed using transposon-mediated approaches combined with next-generation sequencing (e.g. Tn-Seq and TraDIS) (5). Indeed, Tn-seq has identified putative essential genes for *H. influenzae* adaptation to changes in environmental CO_2_ levels, and for survival in the presence of neutrophils, serum and complement (6–8). However, transposon mutagenesis methods are less appropriate for gene essentiality studies as insertion mutants of essential genes are per definition counter-selected. In contrast, inducible Clustered Regularly Interspaced Short Palindromic Repeats interference (CRISPRi) technology, developed from natural CRISPR systems found in bacteria and archaea as part of their adaptive immunity against phages, overcomes this limitation as it enables the transient repression of target genes and allows to monitor phenotypes associated to dispensable and essential genes (9–11). CRISPRi is based on the co-expression of a single-guide RNA (sgRNA) and a catalytically inactive (dead) *Streptococcus pyogenes* Cas9 (dCas9) protein, which binds DNA but lacks endonuclease activity. The sgRNA consists of a 20 nucleotide-long variable spacer sequence that is complementary to a target DNA stretch, a Cas9 handle crucial for interaction with dCas9, and a transcriptional terminator. The sgRNA targets the dCas9 protein to a site on the genome with a sequence complementary to its spacer sequence next to a protospacer adjacent motif (PAM) site composed of 3 nucleotides, typically NGG. When bound to the non-template strand of the target gene, the dCas9•sgRNA complex serves as a roadblock for RNA polymerase, hampering transcription initiation or elongation of the targeted gene (9–14). sgRNAs can be designed to target any sequence of interest provided that the target is next to a PAM site. As such, almost all genes of a given genome can be studied with an appropriate design of sgRNAs if dCas9 and/or the sgRNA is under the control of an inducible promoter to prevent constitutive gene repression. This makes CRISPRi particularly valuable for functional studies including essential gene assessment (15–18).

Two approaches can be employed for genome-wide CRISPRi screenings. The arrayed library method involves handling each mutant with a single sgRNA individually, enabling direct observation of diverse phenotypes (14, 16, 19). This method is laborious and time consuming considering each sgRNA mutant strain needs individual processing. In contrast, the pooled library approach, where multiple knockdown strains are grown together, is more convenient but it limits phenotype measurements besides gene fitness. Deep sequencing reveals changes in sgRNA abundancies of the pooled library upon selective pressures, reflecting the differential fitness provoked by conditional target repression. Hence, the combination of pooled CRISPRi libraries with NGS (CRISPRi-seq) offers a robust method to study gene fitness in bacteria (via relative quantification and comparison of sgRNA abundance) at a genome-wide scale in particular growth conditions (15, 17–26).

Here, we implemented a CRISPRi system suitable for individual genes or pooled libraries knockdown in *H. influenzae*. We first validated operability and titratability of the newly implemented anhydrotetracycline (aTc)-inducible CRISPRi system. Furthermore, we constructed a genome-wide CRISPRi library with high coverage in the genome of RdKW20 strain, designed to cover 99.27% of all total genetic features. As a proof of concept, the generated *H. influenzae* genome-wide CRISPRi library was screened in two laboratory growth media to quantify gene fitness through CRISPRi-seq. We uncovered growth medium-dependent fitness cost, and confirmed several hits with individual knockdown/knockout mutants. In addition, we introduce HaemoBrowse (https://HaemoBrowse.VeeningLab.com), through which the genome annotations and sgRNA design on *H. influenzae* genomes can be readily inspected. The *H. influenzae* CRISPRi platform presented here provides a valuable tool for both gene functional analyses and essentiality in a notorious human pathobiont.

## Results

### An inducible CRISPRi system in *H. influenzae*

In previous work, we successfully established CRISPRi for the Gram-positive human pathogen *Streptococcus pneumoniae* by placing the *dcas9* gene under tight control of LacI and TetR-regulated promoters, while expressing the sgRNA from the synthetic constitutive P3 promoter (16, 18). To adapt this operational streptococcal CRISPRi system for *H. influenzae*, we integrated two cassettes into the chromosome of the widely used laboratory strain *H. influenzae* RdKW20, by taking advantage of its high frequency to naturally acquire exogenous linear DNA. The first cassette was engineered as a linear fragment through Golden Gate cloning encoding *dcas9* under the control of the anhydrotetracycline (aTc)-responsive promoter (hereafter P_tet_) (27), an *erm* erythromycin resistance gene, and the *tetR* transcriptional repressor. The product was integrated into the non-essential *xylB*-*rfaD* locus of the RdKW20 chromosome (28), generating the RdKW20-*dcas9* strain (hereafter, *dcas9* strain) (**Figure 1A**). Integration of the cassette at this locus did not hamper the fitness of the engineered strain compared to its isogenic parental wild-type (**Figure S1A**). In parallel, we constructed the pPEPZHi-*mCherry* vector, which will be used to constitutively express sgRNAs from the P_3_ promoter (13). pPEPZHi-*mCherry* encodes the *mCherry* gene to allow for visual red-white screening of sgRNA cloning in *E. coli*, the dCas9-required handle sequence, and a *spec* spectinomycin resistance gene for selection in both *E. coli* and *H. influenzae*. Illumina read 1 and read 2 sequences flank the sgRNA sequence to allow for a one-step PCR amplification of the sgRNAs and downstream deep sequencing of a library pool. Finally, we introduced flanking homology regions for the neutral Hi0601.1 locus (29), to subsequently generate sgRNA-containing linear cassettes for chromosomal integration in the *dcas9* strain (double homologous recombination) by natural transformation (**Figure 1A**). To assess the efficacy of our CRISPRi system in *H. influenzae*, we cloned three sgRNAs targeting either essential or non-essential (control) genes (30–32) into the chromosome of the P_tet_-*dcas9* background strain. As shown in **Figure 1B**, strains carrying a sgRNA targeting *fabH* (encoding a β-ketoacyl-acyl carrier protein synthase III) or *glyA* (encoding a serine hydroxymethyltransferase) were severely hampered to grow upon addition of 50 ng/mL aTc, in line with their predicted essential functions in the cell. In contrast, the strain with a sgRNA targeting the dispensable gene *sspA* (encoding stringent starvation protein A) grew similarly whether dCas9 was induced or not. As a control, we verified that expression of dCas9 had no effect on growth in a sgRNA-devoid strain (**Figure 1B**, left panel).

**Figure 1.**
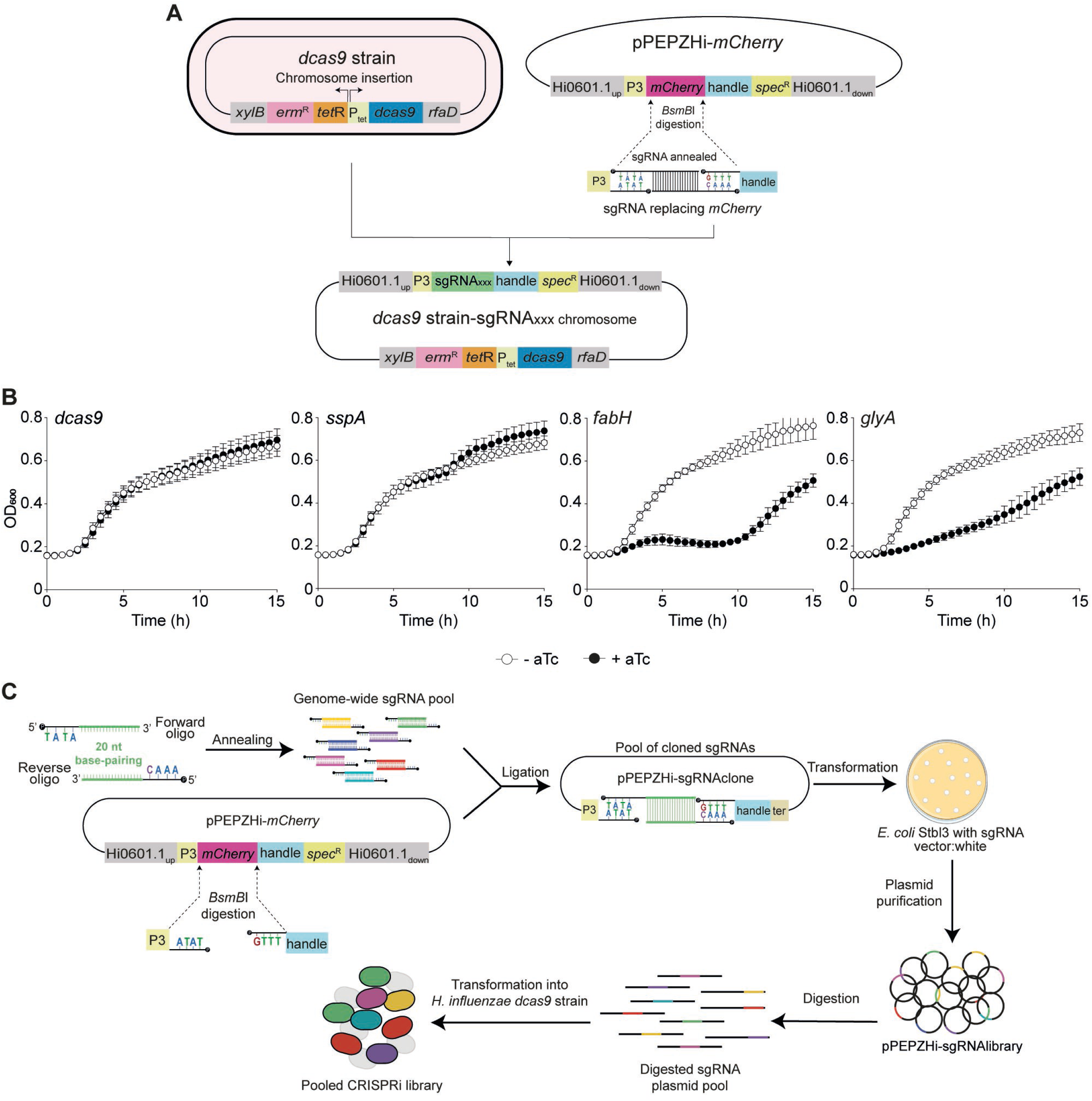
Engineering of a CRISPRi knockdown platform for *H. influenzae*. **(A)** Generation of a *H. influenzae* strain with an aTc-inducible *dcas9* by double homologous recombination at the *xylB*/*rfaD* locus. Backbone vector pPEPZHi-*mCherry* for sgRNA cloning includes a Spec^R^ marker, a P3 promoter controlling sgRNA expression, and an *mCherry* gene encoding a red fluorescent protein flanked by *BsmB*I restriction sites. *BsmB*I vector digestion generates two ends compatible for annealed sgRNA cloning by *mCherry* gene replacement. The sgRNA cassette is amplified from pPEPZHi-sgRNA_XXX_ to perform chromosomal integration at the HI0601.1 *locus* of the *dcas9* background. **(B)** Functional validation of the *H. influenzae* CRISPRi genetic platform. The *dcas9* strain (-) and derivatives expressing *sspA*, *fabH* or *glyA* sgRNAs were grown in sBHI, in the absence (white circles)/presence (black circles) of aTc for CRISPRi-based gene silencing. Growth was monitored by measuring OD_600_ every 30 min for 15 h; standard deviation to the mean is shown for each timepoint. **(C)** Workflow to generate a CRISPRi-based genome-wide library in *H. influenzae*. Oligo pairs containing 20 bp complementary stretchs and 4 nt overhangs compatible with the *BsmB*I-digested pPEPZHi-*mCherry* vector were annealed, phosphorylated and ligated in pool to the *BsmB*I-digested pPEPZHi-*mCherry*. The pool of purified sgRNA-containing plasmids (pPEPZHi-sgRNAlibrary) serves as the reservoir for CRISPRi library construction in *H. influenzae*. The sgRNA library plasmid pool was linearized and transformed into the competent *dcas9* background, generating the pooled CRISPRi library. Both *dcas9* and sgRNA cassettes are integrated in two separate locations into the recipient host chromosome, *xylD/rfaD* and HI0601.1 loci, respectively.

For more subtle control over gene expression, and to determine the dynamic range of the CRISPRi system in *H. influenzae*, we tested the growth profiles for a range of aTc concentrations. We observed a narrow window of gene expression control between 0.25 and 1 ng/mL aTc indicating that the here-used P_tet_ promoter offers some levels of titratability in dCas9 expression (**Figure S1B**). For the rest of the work presented here, we used a saturating aTc concentration of 50 ng/mL.

The results presented demonstrated that the developed CRISPRi system for *H. influenzae* is both functional and inducible. To further verify the accuracy of the system, we selected *ftsZ* and *dnaA* as knockdown targets. Since these targets are directly involved in cell division and DNA replication initiation, respectively, we can evaluate the knockdown phenotype by observing bacterial cell and nucleoid morphology. We engineered sgRNA*_ftsZ_* and sgRNA*_dnaA_* strains as previously described. As expected, cell growth was inhibited for both depletion strains upon dCas9 induction (**Figure 2A**). This macroscopic behavior correlates with an elongation phenotype of most bacterial cells that does not occur when using a sgRNA targeting a neutral gene (*sspA*) (**Figure 2B**). As highlighted by DAPI staining, activation of the CRISPRi system revealed two different phenotypes for *ftsZ* and *dnaA* respective gene knockdowns. Indeed, *ftsZ* knockdown produced filamentous cells with multiple compartmentalized nucleoids or nucleoids that spread throughout the cytoplasm, while cells depleted for *dnaA* harbored a unique compact nucleoid with DNA-free area and produced anucleate minicells (**Figure 2C**). These observations are in direct line with the functions performed by FtszZ and DnaA, demonstrating that the here-described CRISPRi system for *H. influenzae* is specific.

**Figure 2.**
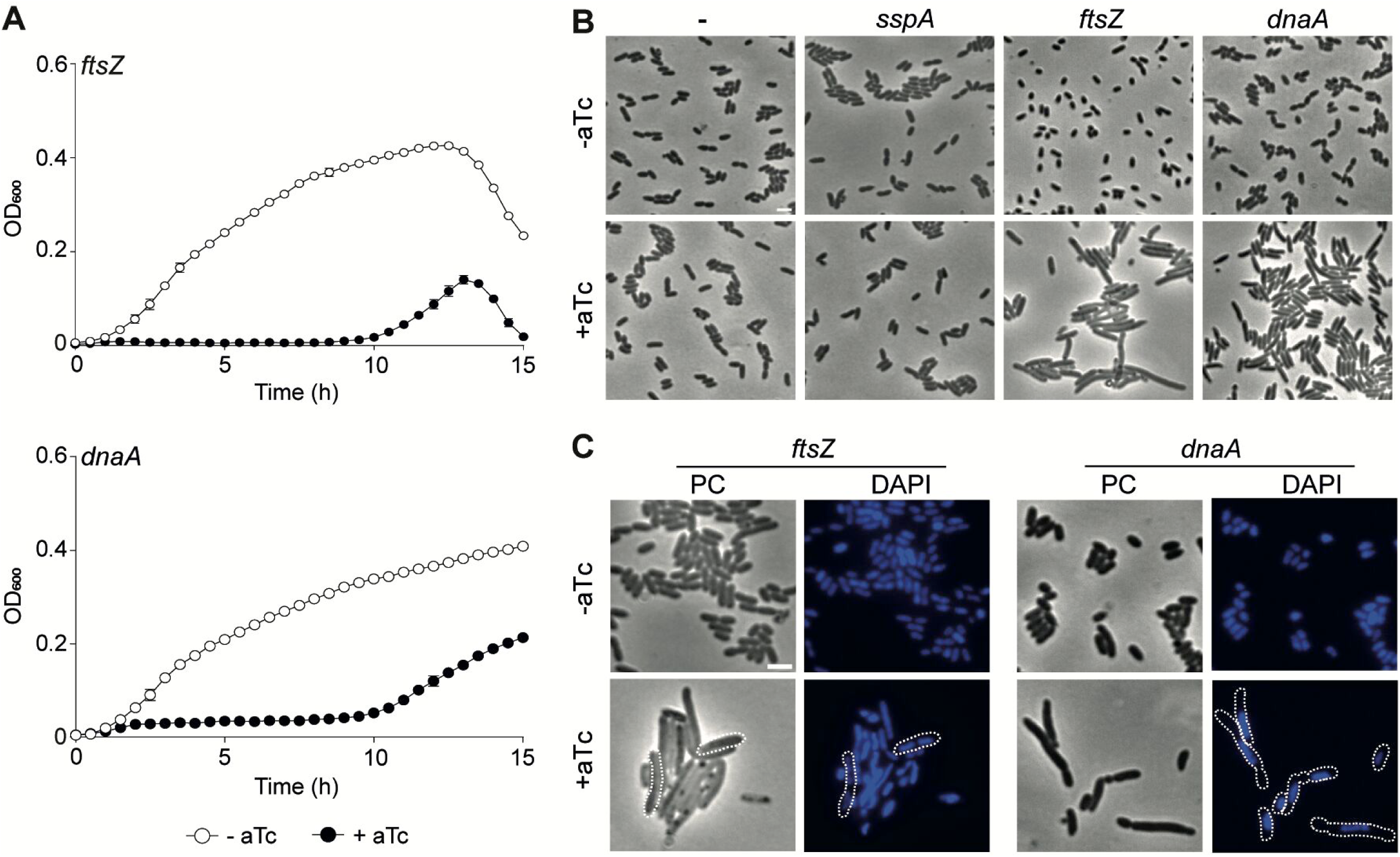
Efficiency and specificity of the *H. influenzae* CRISPRi system. **(A)** The *dcas9* derivative strains expressing *ftsZ* or *dnaA* sgRNAs were grown in sBHI, in the absence (white circles)/presence (black circles) of aTc, in 96-well plates; OD_600_ was measured every 30 min for 15 h. Standard deviation to the mean is shown for each timepoint. **(B)** Phase contrast (PC) microscopy showing bacterial cell morphology for the *dcas9* strains (-) and derivatives expressing *sspA*, *ftsZ* or *dnaA* sgRNAs. Bacteria were grown in sBHI in the absence/presence of aTc. **(C)** Detailed bacterial cell and nucleoid morphology alterations showing the effect of dCas9 induction in *dcas9* derivative strains expressing *ftsZ* or *dnaA* sgRNAs. The nucleoid was stained with DAPI and several cells are outlined to emphasize nucleoid structures and organization in the cytoplasm. Scale bars are equal to 2 µm.

### Development of a genome-wide *H. influenzae* CRISPRi library

To widen the scope of this new tool in *H. influenzae*, we developed a pooled library for strain RdKW20. Through previously described and validated computational algorithms (18), we designed an optimized library including 1,773 sgRNAs targeting all predicted genetic features annotated in the RdKW20 genome. This library covers 99.27% of all total features of the RdKW20 genome (13 features are not targeted by the library due to the lack of PAM sites in their coding sequences). Of note, some designed sgRNAs may target more than one genetic feature if that feature is present in multiple copies, or if repetitive regions are present (**Dataset S1**).

To construct the CRISPRi sgRNA library, we used a synthetic pool of 3,546 ssDNA oligonucleotides for RdKW20 encoding two partially complementary oligonucleotides for each sgRNA (1,773 sgRNAs). From this synthetic ssDNA pool, a pooled plasmid library was constructed through cloning into *BsmB*I-digested pPEPZHi-*mCherry* and transformation into *E. coli* Stbl3 (**Figure 1C**). In total, the cloning yielded ∼10^10^ CFU/mL, ∼4.48 × 10^6^ times the theoretical coverage of the library, of which ∼0.1% was positive for mCherry, indicating a highly efficient digestion and cloning reaction. Subsequent Illumina sequencing of the plasmid library showed a normal distribution of the relative sgRNA prevalence, with only few sgRNAs absent (15) in the pool (**Figure S2A**, **Dataset S2**). The created plasmid library was subsequently used for digestion (see Methods section) and natural transformation of the sgRNA-containing linear cassette pool into the P_tet_-*dcas9* background strain, resulting in ∼1×10^6^ total CFU transformants, and retention of 1,742 sgRNA in a normally distributed library (98.3% of all 1,773 synthesized sgRNAs; 1,701 genes (HI_XXX) targeted by sgRNAs out of 1,779) (**Figure 1C** & **S2B**, **Dataset S2**).

In summary, we were able to generate a CRISPRi library in *H. influenzae* that includes evenly distributed sgRNAs targeting 98.3 % of all annotated features.

### Genome-wide *H. influenzae* fitness evaluation by CRISPRi-seq

Next, we used the CRISPRi library to investigate the RdKW20 strain ‘essentialome’, by quantifying the fitness of each sgRNA target when grown in sBHI medium. For a better resolution of CRISPRi-seq across all sgRNAs, three time points (7, 14 and 21 generations) were sampled in quadruplicate (**Figure 3A**). Induction of dCas9 was achieved by adding aTc at 50 ng/mL.

**Figure 3.**
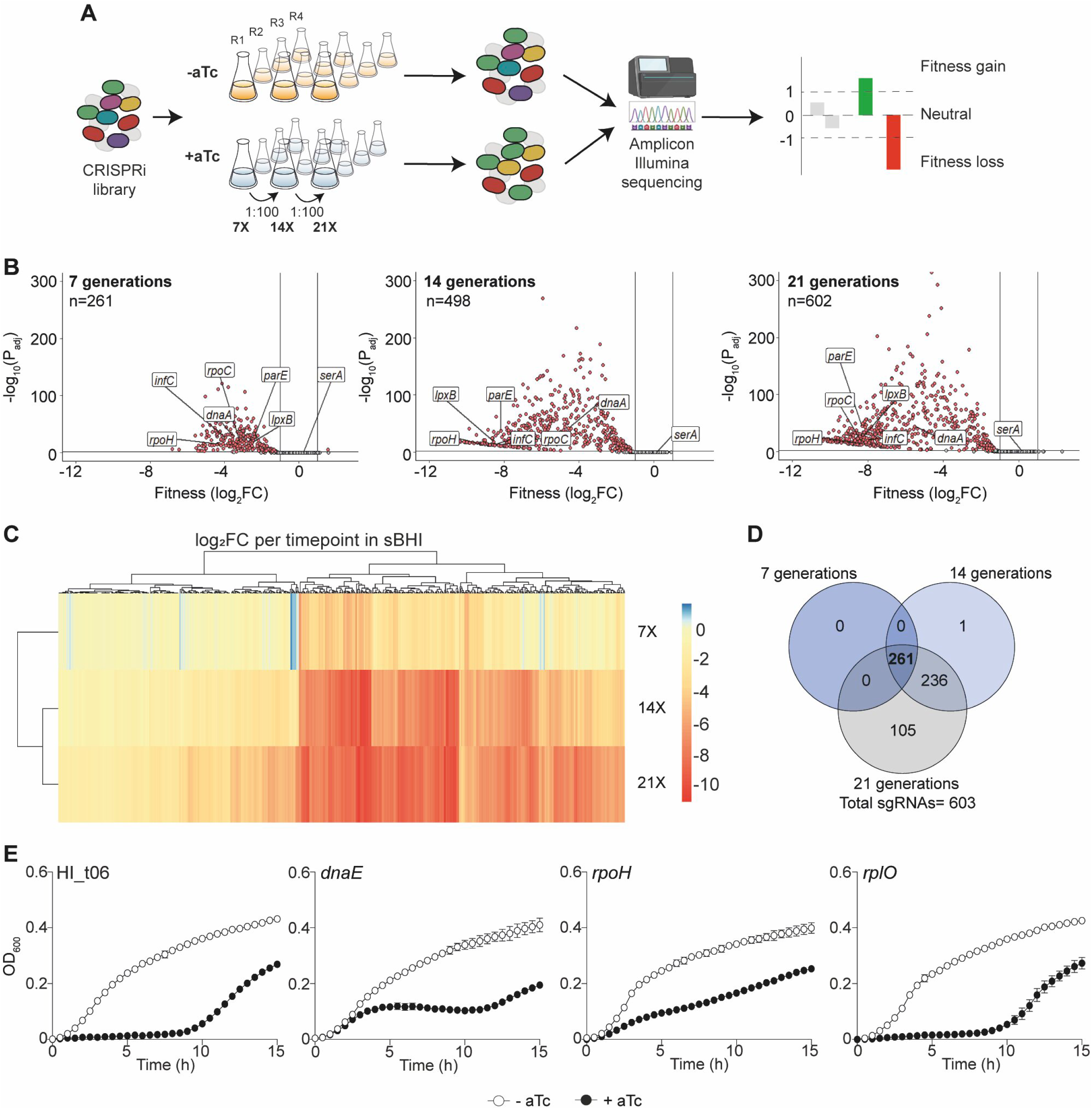
CRISPRi-seq gene fitness analysis of *H. influenzae* during growth in sBHI medium. **(A)** The *dcas9* CRISPRi library was grown in sBHI without (orange) or with (blue) aTc 50 ng/mL. Four biological replicates were performed for each condition. After 7, 14 and 21 generations of exponential growth, bacterial cultures were pelleted, genomic DNA was purified, sgRNA-containing amplicons were generated and Illumina sequenced, and data were analysed for differential fitness score abundance. **(B)** Volcano plots show sgRNA target fitness scores upon *dcas9* induction for 7, 14 and 21 generations of exponential growth in sBHI. Red dots represent sgRNAs targeting essential genes, with significant differential fitness effects (log_2_FC<−1 and P_adj_<0.05; vertical and horizontal black lines). Several essential (*dnaA* [DNA replication initiation], *infC* [translation initiation factor], *lpxB* [lipid A synthase], *parE* [DNA topoisomerase], *rpoC* [RNA polymerase] and *rpoH* [sigma factor]) and dispensable (*serA* [serine metabolism]) genes in sBHI have been plotted. **(C)** The heatmap combined with hierarchical clustering displays all significantly enriched/depleted sgRNAs over the three timepoints. A negative (red) score indicates a fitness loss in inducing conditions, while a positive (blue) score indicates a fitness gain. **(D)** Venn diagram showing the overlap of sgRNAs with reduced abundance after aTc induction over the three timepoints. **(E)** CRISPRi-based analysis of *H. influenzae* specific gene fitness. *dcas9* derivative strains expressing HI_t06*, dnaE*, *rpoH* or *rplO* sgRNAs were grown in sBHI, in the absence (white circles)/presence (black circles) of aTc. Strains were grown in 96-well plates, and OD_600_ was measured every 30 min for 15 h. Standard deviation to the mean is shown for each timepoint.

Through deep sequencing, quantification and comparison of sgRNA abundance in induced *versus* uninduced samples, we confirmed the tight control over *dcas9* expression by P_tet_, as all uninduced samples had a similar sgRNA composition, whether they were grown for 7, 14, or 21 generations (**Figure S3A & B**). Considering induced samples, the majority (91%) of variance between samples was due to *dcas9* induction in a time-dependent manner. Fitness score comparison between the three conditions showed that the number of essential genes gradually increased according to the number of generations (**Figure 3B, C & D**, **Dataset S3**). Indeed, within the induced samples, we observed that 261 sgRNAs have significantly reduced abundance after 7 generations, 498 after 14 generations and 602 after 21 generations (**Figure 3B & D**, **Dataset S3**). Consistently, sgRNAs with significant lower abundances at earlier timepoints remain so at later timepoints (**Figure 3D**). Moreover, volcano plots and heatmaps highlight the trend for individual fitness scores to be lower and lower over generations (**Figure 3B & C**, **Figure S3B**). In direct line with individual sgRNA data (**Figure 1B & 2**), sgRNAs targeting *glyA, dnaA, fabH* or *ftsZ* showed drastic reduced abundances (**Dataset S3**). To validate this CRISPRi-seq fitness screen performed in sBHI growth medium, we independently analyzed a panel of genes whose sgRNAs showed significantly reduced abundance after 7 generations. Fourteen separate sgRNAs targeting the HI_t06, *dnaE*, *rpoH*, *rplO*, *infC*, *rpsL*, *ispE*, *metK*, *rpoC*, *lpxB*, *parE*, *rsxA, rpsT* and *mepA* genes were individually cloned in the pPEPZHi-*mCherry* vector and integrated in the P_tet_-*dcas9* background, as described above. In the presence of inducer, twelve out of the fourteen tested sgRNAs strains exhibited growth defects (**Figure 3E & S4**).

These results strongly validate the robustness and reliability of this novel *H. influenzae* CRISPRi platform in bacterial genome-wide assays, and unveil gene requirements in a laboratory-based complex medium.

### *H. influenzae* CRISPRi-seq reveals growth medium-dependent essentiality

We next assessed *H. influenzae* gene essentiality in a different growth condition through cultivation in a chemically defined (CDM) medium (33, 34), and subsequent comparative analysis to the sBHI data. The CRISPRi library was grown in CDM in the absence/presence of aTc, and deep-sequenced for gene fitness quantification. Akin to sBHI, variability was primarily attributed to *dcas9* expression with longer induction times increasing differences in sgRNA content (**Figure S3A & B)**, as the number of genes with significant fitness defects were 259 sgRNAs with reduced abundance after 7; 513 after 14; and 590 after 21 generations (**Figure 4A, B & C**, **Dataset S3**). As reported hereabove for the sBHI screen, we validated the functionality of this high-throughput CRISPRi screen in CDM with 12 (out of 14) individual sgRNA strains (**Figure 4D & S5**).

**Figure 4.**
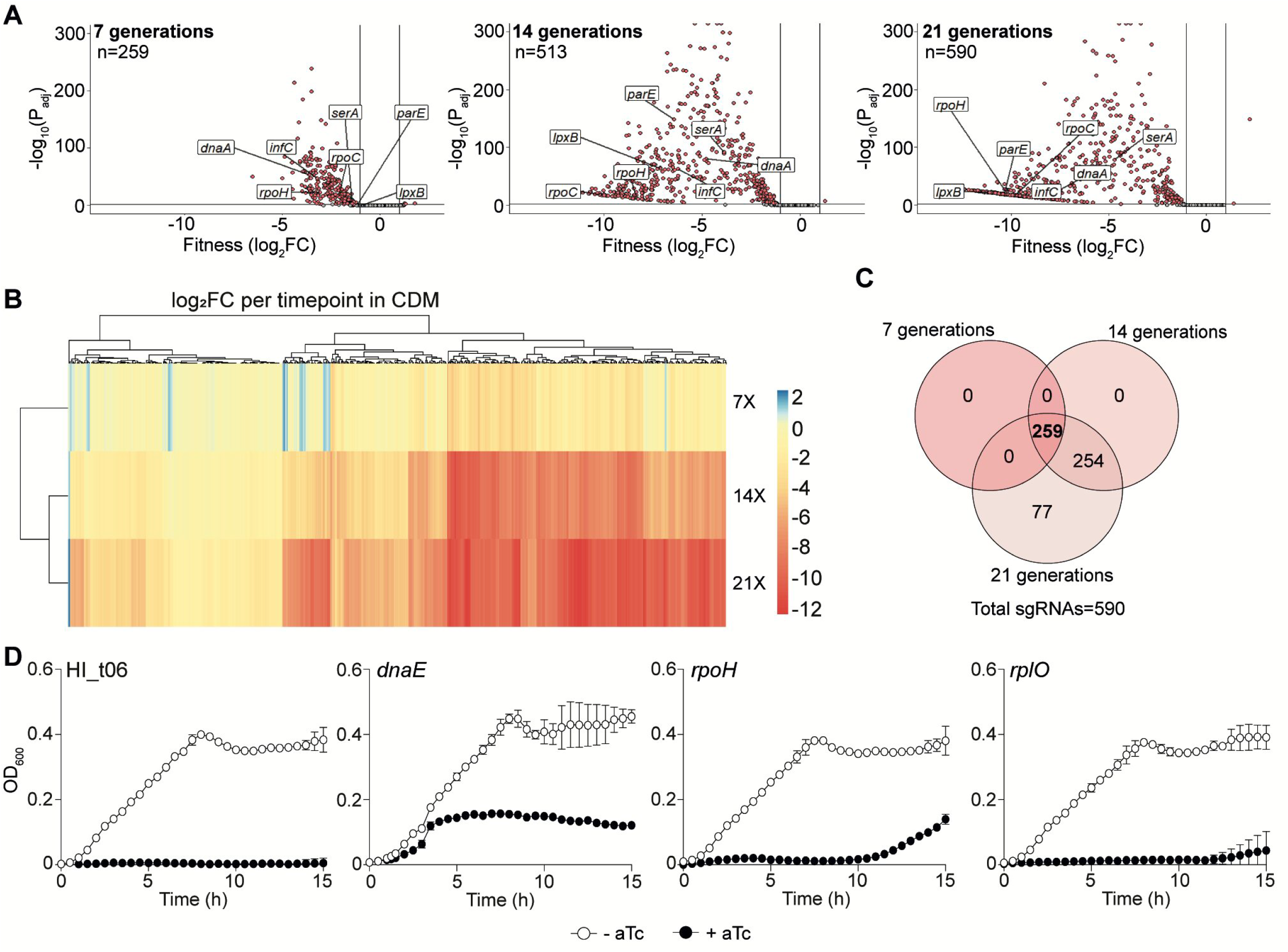
CRISPRi-seq gene fitness analysis of *H. influenzae* during growth in CDM medium. **(A)** Volcano plots show sgRNA target fitness scores upon *dcas9* induction for 7, 14 and 21 generations of exponential growth in CDM. Red dots represent sgRNAs targeting essential genes, with significant differential fitness effects (log_2_FC<−1 and P_adj_<0.05; vertical and horizontal black lines). Several essential genes (*dnaA* [DNA replication initiation], *infC* [translation initiation factor], *lpxB* [lipid A synthase], *parE* [DNA topoisomerase], *rpoC* [RNA polymerase], *rpoH* [sigma factor] and *serA* [serine metabolism]) in CDM have been plotted. **(B)** The heatmap combined with hierarchical clustering displays all significantly enriched/depleted sgRNAs over the three timepoints. A negative (red) score indicates at fitness loss in inducing conditions, while a positive (blue) score indicates a fitness gain. **(C)** Venn diagram showing the overlap of sgRNAs with reduced abundance after induction over the three timepoints. **(D)** *dcas9* derivative strains expressing HI_t06*, dnaE*, *rpoH* or *rplO* sgRNAs were grown in CDM, in the absence (white circles)/presence (black circles) of aTc. Strains were grown in 96-well plates, OD_600_ was measured every 30 min for 15 h. Standard deviation to the mean is shown for each timepoint.

A significant number of essential genes were common to both sBHI and CDM media (531 genes). As expected, analysis of Clusters of Orthologous Genes (COG) for all essential genes (80.3% shared essential genes between the two media) showed minor differences. Translation, ribosomal structure and biogenesis was the most affected category. Small differences between the two media were observed for inorganic ion transport, nucleotide and amino acid metabolism and transport, as well as energy production (**Figure 5A**).

**Figure 5.**
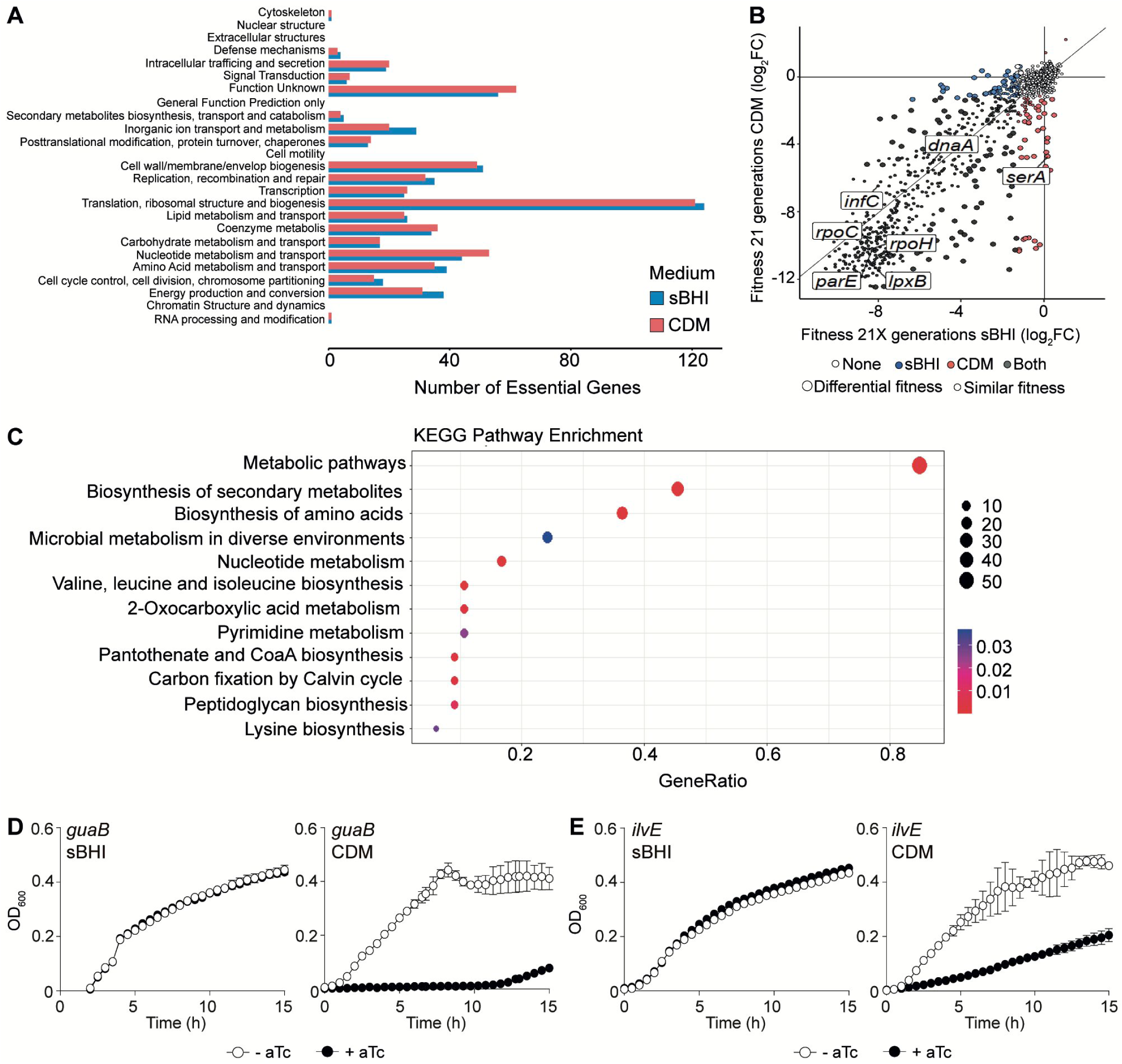
CRISPRi-seq reveals medium-specific growth determinants for *H. influenzae*. **(A)** Distribution of COG categories in all significantly essential genes in sBHI (blue) and CDM (red). **(B)** Interaction plot comparing fitness scores upon *dcas9* induction for 21 generations in sBHI *versus* CDM. Blue and red dots represent sgRNAs with log_2_FC<−1 and P_adj_<0.05 in sBHI and CDM, respectively. Black dots represent sgRNAs with log_2_FC<−1 and P_adj_<0.05 in both conditions, and white dots color dispensable genes. Dot size refers to genes considered as differentially essential in one medium (94 genes more essential in CDM; 60 genes more essential in sBHI). Several genes (*dnaA* [DNA replication initiation], *infC* [translation initiation factor], *lpxB* [lipid A synthase], *parE* [DNA topoisomerase], *rpoC* [RNA polymerase], *rpoH* [sigma factor] and *serA* [serine metabolism]) essential in at least one medium are labeled. **(C)** KEGG pathway enrichment analysis in CDM (*versus* sBHI). The gene ratio refers to the number of enriched genes in CDM for a specific pathway reported to the total amount of differentially essential genes. Dot size reflects the number of differentially essential genes. Color shades scale the significance (P_adj_). **(D & E)** *dcas9* derivative strains expressing sgRNA*_guaB_* (**D**) or sgRNA*_ilvE_* (**E**) were grown in sBHI and CDM, in the absence (white circles)/presence (black circles) of aTc. Strains were grown in 96-well plates, OD_600_ was measured every 30 min for 15 h, standard deviation to the mean is shown for each timepoint.

Next, we performed a differential fitness analysis between samples grown in sBHI *versus* CDM. Comparative fitness analysis showed that 7, 35 and 60 genes were significantly more essential in sBHI at 7, 14, and 21 generations, respectively, whilst 22, 75 and 94 genes were significantly more essential in CDM (**Figure 5B**, **Datasets S4** & **S5**). Through a KEGG pathway enrichment analysis on the genes differentially essential after 21 generations, we observed that more essential genes in CDM mostly belong to amino acid and nucleotide biosynthesis pathways (36% and 17%, respectively) and metabolic pathways in a more general manner (85%) (**Figure 5C**). In contrast, sBHI-specific essential genes primarily encode transporters (35%) (**Figure S6**). Aiming to confirm this medium-dependent essentiality, we selected 2 genes, *guaB* and *ilvE*, whose sgRNAs showed reduced abundance in CDM but stayed equal in sBHI. The *guaB* gene encodes an inosine-5’-monophosphate dehydrogenase that catalyzes the conversion of inosine 5’-phosphate to xanthosine 5’-phosphate, the first committed and rate-limiting step in the *de novo* synthesis of guanine nucleotides. The *ilvE* gene encodes for a branched chain amino acid aminotransferase acting on leucine, isoleucine and valine. Both sgRNAs were individually cloned in the P_tet_-*dcas9* background and bacterial growth was assayed in CDM and sBHI (in the absence/presence of aTc). Strikingly, dCas9 induction showed a CDM-specific growth defect, while growth in sBHI was unaffected regardless of dCas9 induction (**Figure 5D & E**).

These results validate the use of this CRISPRi platform in bacterial genome-wide assays to reveal growth condition-dependent gene requirements.

### CRISPRi-programmed fitness defects are rescuable in *H. influenzae*

Our differential fitness analysis between sBHI or CDM grown samples shed light on medium specificities and genetic targets (**Figure S7** & **S8**) (30, 35). Among others, genes encoding enzymes required for threonine and serine biosynthesis are differentially essential in CDM compared to sBHI (**Figure 6 & S9**). As a last validation of this *H. influenzae* CRISPRi system, we tested whether the observed CRISPRi-promoted fitness burden could be rescued by other genetic tools. We selected the CDM-specific *serA* gene, which encodes a D-3-phosphoglycerate dehydrogenase involved in the L-serine biosynthesis pathway. Therefore, repression of the *serA* gene should cause L-serine auxotrophy. Coherently, in the presence of aTc, bacterial growth of the sgRNA*_serA_* strain exhibited a severe defect in CDM, but not in sBHI (**Figure 6A**). In line with this data, a Δ*serA* deletion mutant showed auxotrophy when grown in CDM (**Figure 6B**). L-serine concentration is about 285 mM in sBHI (Javier Asensio-López, personal communication), while only 0.29 mM in CDM (35). We reasoned that increasing L-serine concentration in CDM might abolish the sgRNA*_serA_* and Δ*serA* fitness defects. As expected, supplementation of CDM with L-serine 10 mM abolished the growth arrest phenotype caused by underexpression of *serA* (**Figure 6B & C**). For completion, similar observations were made for the *dcas9*-sgRNA*_serA_* and Δ*serA* strains when using a minimal chemically defined medium deprived of L-serine (mCDM) (**Figure S10**). These results fully support the operativity of our study design and CRISPRi platform to identify *H. influenzae* genes undergoing medium-dependent fitness cost.

**Figure 6.**
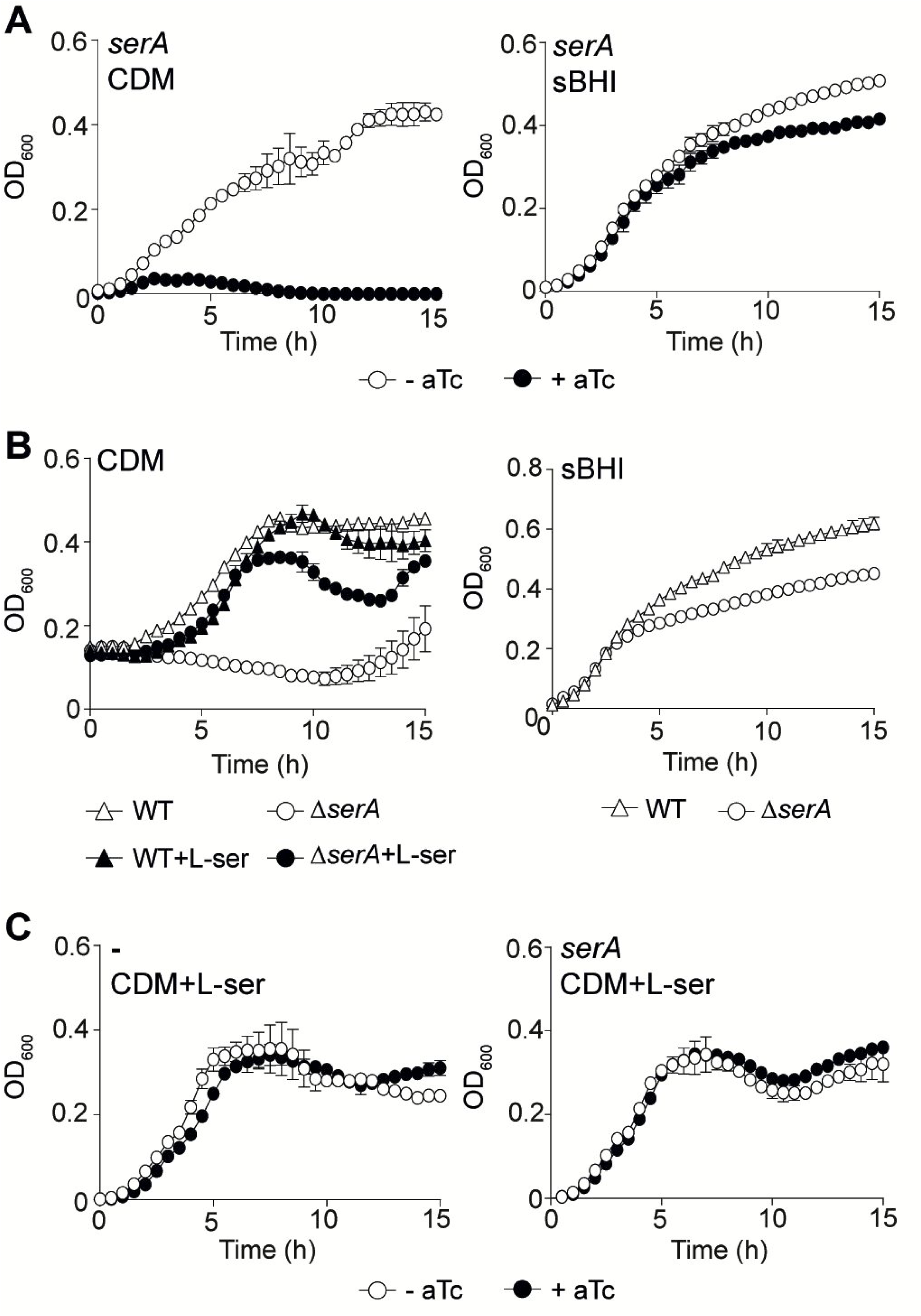
sgRNA gene silencing reversibility in *H. influenzae*. **(A)** The *dcas9* derivative strain expressing sgRNA*_serA_*was grown in CDM (left panel) and sBHI (right panel), in the absence (white circles)/presence (black circles) of aTc. **(B)** RdKW20 WT (triangles) and Δ*serA* (circles) strains were grown in CDM in the absence (white) or presence (black) of L-serine 10 mM (left panel), and in sBHI (right panel). **(C)** *dcas9* strain and its derivative expressing sgRNA*_serA_* were grown in CDM supplemented with L-serine 10 mM, in the absence (white circles)/presence (black circles) of aTc. Strains were grown in 96-well plates, OD_600_ was measured every 30 min for 15 h, standard deviation to the mean is shown for each timepoint.

### HaemoBrowse: a visual and intuitive *H. influenzae* genome browser

Finally, we developed HaemoBrowse, a genome browser for *H. influenzae* (**Figure 7**). Based on JBrowse 2, HaemoBrowse offers a user-friendly and highly intuitive visual platform to browse the genome of the strain RdKW20 for its encoded features. Moreover, HaemoBrowse shows the position and sequence of the designed sgRNAs designed. Additional flexibility is offered by allowing for the search of locus tags (including HI_XXXX, locus tags for RdKW20) and gene names. We also included the genomes of three other commonly used *H. influenzae* strains, NTHi375, R2866 and 86-028NP. To allow for further exploration of gene function, direct links to the UniProt, AlphaFold, FoldSeek, and STRING databases are available for each encoded feature. Finally, the conservation and the essentiality of the encoded features in the different conditions and at different timepoints, as determined through the CRISPRi-system introduced in RdKW20 here, are documented. HaemoBrowse is freely available at https://HaemoBrowse.VeeningLab.com.

**Figure 7.**
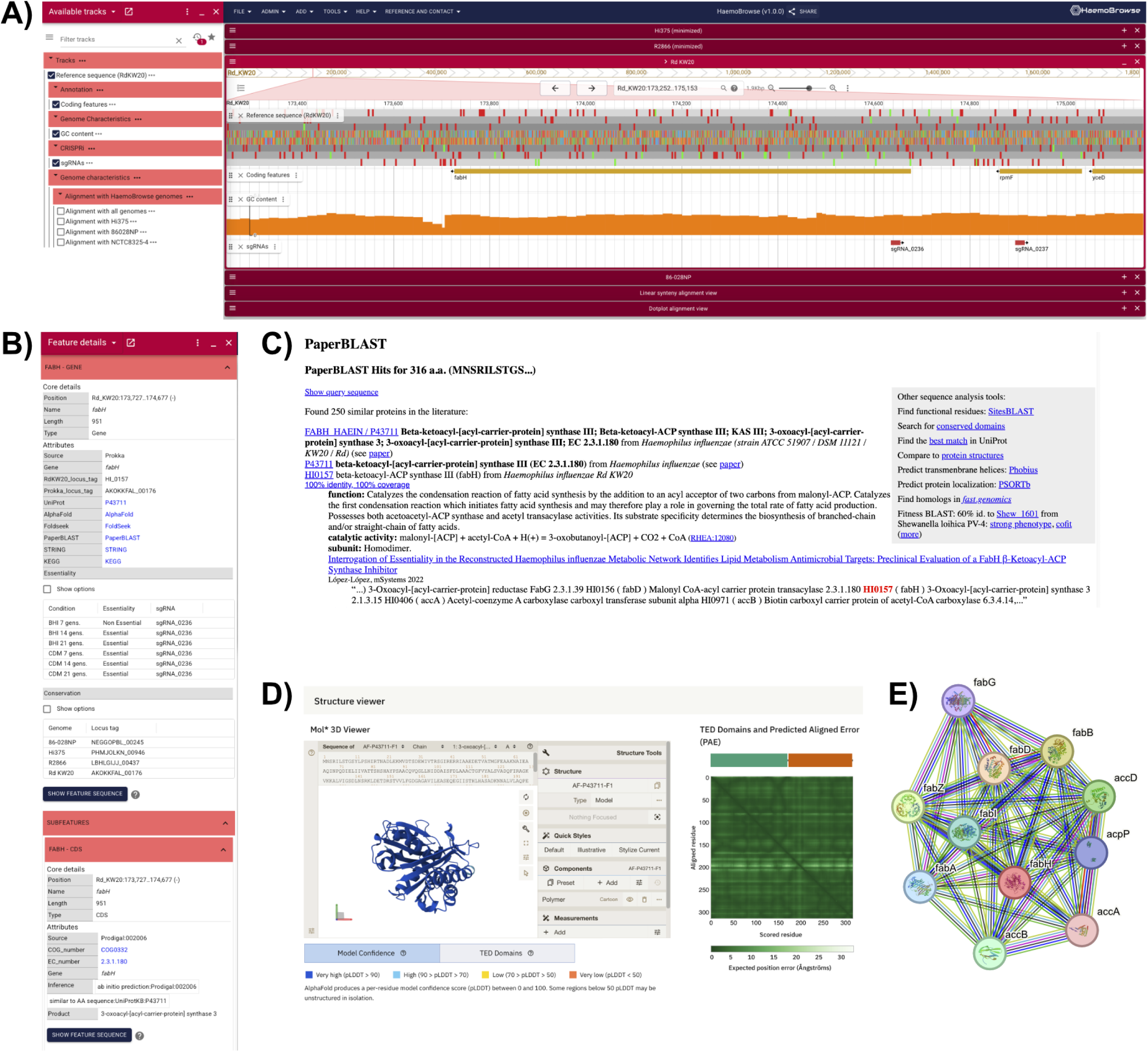
HaemoBrowse, available at https://HaemoBrowse.VeeningLab.com. **(A)** A screenshot of the *fabH locus* as shown in HaemoBrowse. In the left panel, tracks can be turned on/off. In the right panel, the genome can be browsed by dragging the mouse to the left or right, zooming in and out, or searched on gene name and/or *locus* tags. **(B)** For each coding sequence, a context menu provides additional information and links to external resources, such as PaperBLAST, AlphaFold, and STRING. The context menu also displays the essentiality per experimental condition. (**C**) PaperBLAST provides additional information through literature search. (**D**) Structure predictions can be consulted through AlphaFold. (**E**) STRING database shows interactions of the protein of interest.

## Discussion

Identification of bacterial genetic targets for drug development requires powerful screening strategies and extended information on gene function and bacterial physiology to lead such pharmacological search. Here, we made a step forward towards this goal for *H. influenzae* by designing a robust genome-wide screen. We engineered a scalable CRISPRi platform and developed a CRISPRi library covering 99.27% of all annotated features in the genome of the *H. influenzae* RdKW20 strain. We demonstrate the functionality of our CRISPRi system at a macroscopic and microscopic scale in the RdKW20 strain, and employed CRISPRi-seq to quantify bacterial gene fitness *in vitro* on a genome-wide level. Moreover, given that this CRISPRi system is responsive to any type of tetracycline-like molecule including doxycycline, challenging vertebrate/murine models with an induced/uninduced pool of our CRISPRi library is likely to identify genes involved in virulence, invasion or colonization at multiple steps of infection, as demonstrated for *S. pneumoniae* (18).

Our high-throughput knockdown approach is validated by a previous transposon-based knockout essentialome study in sBHI (36). Even more, 87.3% (219 out 251) of the essential genes identified via a transposon-mutant library across independent studies (36-37 & Gil-Campillo, personal communication) were also identified at 21 generations in sBHI. Importantly, the inducibility of our system allowed us to refine the study of essential genes. For instance, 47 genes that are significantly essential in both CDM and sBHI are considered as more essential in CDM, providing an exhaustive view of the genetic requirements for bacterial growth in this specific medium. This kind of differential analysis on crucial genes is obviously precluded with Tn libraries where clones bearing a Tn insertion in an essential gene are counterselected and eradicated from the pool of Tn mutants. Also, sampling at multiple time points provided a dynamic view of *H. influenzae* genes important for fitness. In a logical way, a prolonged inducer treatment aggravates fitness losses, resulting in larger fold changes. This may also help to identify genes with marginal fitness effects over generations. In addition, this temporal analysis gives information about the kinetics of depletion that, besides the efficiency of sgRNA repression, reflects the necessity, abundancy and stability of the encoded protein.

Of note, a decrease in the abundance of an sgRNA does not automatically indicate that the gene is essential as such, but rather less competitive than the uninduced control under a given condition. Therefore, CRISPRi data interpretation strongly requires considering the robustness of the significance. In addition, CRISPRi is by nature polar thus multiple genes in polycistronic operons can be affected by a single sgRNA. It is thus always advised to consider the genetic context of the CRISPRi knockdown, something that is now facilitated by the generation of HaemoBrowse (**Figure 7**). Moreover, the pooled growth approach described here is robust and convenient, but unable to differentiate between competitive and environmental influences on sgRNA selection, as sgRNA selection in the sample may be influenced when cells with different sgRNA content grow together. Thus, as it is shown in **Figures 3, 4, S4 & S5**, it is required to perform confirmation experiments. Notably, our confirmation assays strongly validate the robustness and reliability of the CRISPRi-seq platform presented in this study.

Importantly, our differential essentialome analysis showcased medium specificities. Depletion of many genes involved in purine (*guaB*) and amino acid (e.g. *ilvE* and *serA*) metabolism showed a marked impact on growth in CDM when compared to sBHI. The nutrient scarcity in CDM is supposed to be the main reason for such requirements. Notably, the system may also have the potential to titrate metabolite concentration requirements, as sgRNA*_serA_* abundance drops in a medium containing about 10 times less molecules of L-serine (CDM). Likewise, knockdown defect restoration is feasible, as seen upon addition of exogenous extra L-serine in CDM assays. The differentially essential gene enrichment in sBHI is more puzzling. Depleting ABC transporter systems for metal (Zn, Mo or Ni) or carbon sources (xylose, dipeptides, methionine) in a complex and rich medium such as sBHI is toxic. This might be due to a defect in pumping out nutrients from cytoplasm that provoke an accumulation of toxic intermediates. These observations together reinforce the versatility of our CRISPRi platform, and open a whole range of clinically relevant *in vitro* or *in vivo* conditions to be investigated in future studies.

In summary, the genetic (inducible promoter, neutral insertion platform, depletion system, genome-wide fitness library) and *in silico* (HaemoBrowse) resources generated here provide powerful tools to deeply investigate the molecular genetics of *H. influenzae*, facilitating the understanding of this pathogen physiology and the identification of therapeutic targets.

## Materials and Methods

### Bacterial strains, growth conditions and chemicals

Strains used in this study are listed in **Table S1**. *H. influenzae* strains were grown on solid medium at 37°C with 5% CO_2_ using either PolyVitex agar (PVX, bioMérieux, 43101), or *Haemophilus* Test Medium agar (Oxoid, CM0898) supplemented with 10 μg/mL hemin and 10 μg/mL nicotinamide adenine dinucleotide (NAD) (sHTM). *H. influenzae* liquid cultures were grown at 37°C with 5% CO_2_ in either brain heart infusion medium (Oxoid, CM1135), chemically defined medium (CDM) or minimal chemically defined medium (mCDM, see **Table S2**), in all cases supplemented with 10 μg/mL hemin and 10 μg/mL NAD (referred to sBHI, CDM or mCDM). Stock solutions of anhydrotetracycline 500 µg/mL (aTc, Merck, 37919-100MG-R) were prepared in ethanol 96%. L-Serine (Merck, S4500) 0.47 M stock solution was prepared in distilled water. For *H. influenzae*, erythromycin 11 μg/mL (Erm_11_), spectinomycin 50 μg/mL (Spec_50_), or L-serine 10 mM were used when necessary. *Escherichia coli* was grown on Luria Bertani (LB) or LB agar at 37°C, supplemented with Spec_50_ when appropriate.

### Construction of a *H. influenzae* host strain with an aTc-inducible dCas9

Plasmids and primers used in this work are shown in **Tables S3** and **S4**, respectively. The aTc-inducible system to control *dcas9* expression was based on a previously developed doxycycline-inducible CRISPRi system in *S. pneumoniae* (18). Four fragments (1 to 4) were amplified separately for assembly using Golden Gate cloning. Fragment 1, containing the *H. influenzae xylB* region for homologous recombination, was amplified from RdKW20 genomic DNA with F_*xylB*/2469 and R_*Aar*I_*xylB*-ery/2470 primers; fragment 2, containing an Erm (*ermC*) selection marker, was amplified from pUC19-Hi061.1-Erm with F_*Aar*I_ery/2471 and R_*Aar*I_ery/2472 primers; fragment 3, containing the *tetR*-P_tet_-*dcas9* cassette, was amplified from *S. pneumoniae* VL3468 strain genomic DNA (18) with F_*Aar*I_*tetR*/2473 and R_*Aar*I_dCas9/2474 primers; and fragment 4, containing the *H. influenzae rfaD* region for homologous recombination, was amplified from RdKW20 genomic DNA with F_*Aar*I_*rfaD*/2475 and R_*rfaD*/2476 primers. The four DNA fragments were separately purified using NucleoSpin Gel and PCR Clean-up Kit (Macherey-Nagel, 22740609.250), and assembled by Golden Gate cloning to build the P_tet_-*dcas9* cassette. The assembly reaction, performed with *Aar*I (New England Biolabs, ER1581) and T4 DNA ligase (New England Biolabs, M0202S), involved 30 cycles of 1.5 min/cycle at 37°C, followed by 3 min at 16°C. Enzymes were inactivated at 80°C for 10 min. The 8,424 bp ligation product was amplified by PCR with primers F_*xylB*/2469 and R_*rfaD*/2476, purified using NucleoSpin Gel and PCR Clean-up Kit, and chromosomally integrated in the RdKW20 genome by double homologous recombination between the *xylB* and *rfaD* genes using the MI-V medium (38). Chromosomal integration was selected on sHTM-agar with Erm_11_, to generate the RdKW20 derivative strain RdKW20-*dcas9*, used as a host recipient strain for individual sgRNA and sgRNA library cloning (named as *dcas9* strain).

### Generation of a vector platform for sgRNA cloning

Five fragments were assembled to generate the backbone vector pPEPZHi-*mCherry*, a derivative of pPEPZ-sgRNAclone (15, 18). Fragment 1, containing the pUC18 replication origin, was amplified from pPEPZ-sgRNAclone vector with F_*Aar*I-oUC18/2483 and R_*Aar*I-oUC18/2484 primers; fragment 2, containing the *H. influenzae* Hi0601.1 upstream region for homologous recombination, was amplified from RdKW20 genomic DNA with F_*Aar*I-Hi0601up/2477 and R_*Aar*I-Hi0601up/2478 primers; fragment 3 containing Illumina read 1 sequence, P3 promoter, *mCherry*-encoding gene flanked by *BsmB*I restriction sites, dCas9 handle binding, terminator region of sgRNA, Illumina read 2 sequence, 8 bp Illumina index sequence and P7 adaptor sequence, was amplified from pPEPZ-sgRNAclone with primers F_*AarI*-P3-sgRNA/2488 and R_*Aar*I-sgRNA/2480; fragment 4, containing the 5’ end of a Spec^R^ gene (nucleotides from 1 to 1,333) was amplified from pUC19-Hi061.1-Spec (39) with F_*Aar*I-specHi/2485 and R_*Aar*I_specmut/2517 primers; fragment 5 containing the rest of the same Spec^R^ marker (nucleotides from 1,334 to 1,948) and the *H. influenzae* Hi0601.1 *locus* downstream region for homologous recombination, was amplified from pUC19-Hi061.1-Spec with F_*Aar*I_specmut/2518 and R_*Aar*I_Hi0601dn/2482 primers. The five fragments were purified separately using NucleoSpin Gel and PCR Clean-up Kit, and used in a Golden Gate assembly reaction with *Aar*I and T4 ligase, for 30 cycles of 1.5 min at 37°C followed by 3 min at 16°C. Enzymes were inactivated at 80°C for 10 min. The Golden Gate product, i.e. backbone vector, was directly used to transform chemically competent *E. coli* Stbl3. Growth of red Spec^R^ colonies after plating indicated successful transformation of *E. coli* Stbl3 with pPEPZHi-*mCherry*.

### Construction of individual sgRNA plasmids

Targeting sequences were ordered from Stab Vida (https://www.stabvida.com/es) as two partially complementary 24 nt primers (**Table S4**). Annealing of each oligo pair generated a dsDNA fragment with 20 nt sgRNA base-pairing spacer sequence and 4 nt overhang at each end. The annealing reaction was performed separately for each sgRNA in 10xTEN buffer (10 mM Tris, 1 mM EDTA, 100 mM NaCl, pH 8) in a thermocycler, for 5 min at 95°C, followed by slow cooldown at RT (0.1°C/s). The two 4 nt overhangs were designed to be compatible with the two adhesive ends of the *BsmB*I-digested pPEPZHi-*mCherry*. The pPEPZHi-*mCherry* vector was *BsmB*I*-*digested (Fisher Scientific, ER0451) and purified by gel extraction to ensure removal of the 739 bp *mCherry* gene. Each annealing product was individually ligated into *BsmB*I-digested pPEPZHi-*mCherry* with T4 ligase to generate pPEPZHi-sgRNA*geneX*clone plasmids. Ligation products were transformed into chemically competent Top10 *E. coli* cells. Transformants were selected on LB agar with Spec_50_. After red/white screening and PCR confirmation, one colony from each pPEPZHi-sgRNAclone was selected for culturing and plasmid purification using PureYieldTM Plasmid Miniprep System (Promega, A1222). Purified plasmids were used to amplify each sgRNA cassette by PCR with F_AarI-Hi0601up/2477 and R_AarI_Hi0601dn/2482 primers. PCR products were transformed into the *dcas9* strain, generating RdKW20-*dcas9* P3-sgRNA*_geneX_*. Three individual colonies per construct were stored and used for later analyses.

### Growth assay of *H. influenzae* CRISPRi system with individual sgRNAs

*H. influenzae dcas9* P3-sgRNA*_geneX_* clones were grown on PVX agar for 12 h at 37°C with 5% CO_2_. Two to five colonies were inoculated in 10 mL sBHI or CDM and grown for 12 h with shaking (90 r.p.m). Cultures were then diluted to OD_600_=0.05, incubated in sterile 50 mL flasks with 10 mL sBHI or CDM and shaking (180 r.p.m), up to OD_600_=0.3. Cultures were diluted for a second time to OD_600_=0.03, each in duplicate, incubated in sterile 50 mL flasks with 10 mL sBHI or CDM, in the presence or absence of aTc 50 ng/mL for 2 h at 180 r.p.m, up to OD_600_=0.3. Afterwards, cultures were diluted to OD_600_=0.01 in sBHI or CDM, and 198 µL-aliquots were transferred to individual wells in 96-well plates, in the respective presence or absence of aTc 50 ng/mL. Plates were incubated at 37°C for 15 h in an Agilent Biotek Synergy H1 Microplate Reader H1M Unit3 – AV; OD_600_ was recorded every 30 min. For the P_tet_ responsiveness experiment, cells were grown for 12 h at 37°C in 10 mL sBHI with shaking (120 r.p.m). Next, cultures were diluted in fresh sBHI to OD_600_=0.01 and transferred to a 96-wells plate with a range of aTc concentrations. In all cases, each growth curve was corrected to its respective blank value. All 96-well experiments were performed in duplicate on at least three independent occasions (n≥3).

### sgRNA library design

Prokka (40) was used to annotate the *H. influenzae* genomes RdKW20, NTHi375, 86-028NP and R2866, and the non-redundant sgRNA library was automatically designed via a R-pipeline (https://github.com/veeninglab/CRISPRi-seq) to target every genetic feature as previously described (15, 41). The list of 7,260 sgRNAs designed for the 4 aforementioned strains was narrowed down to generate a non-redundant library of 3,351 sgRNAs (1,773 sgRNAs specific to strain RdKW20) and avoid exact duplicates that will bias the library distribution. Detailed information about the list of all sgRNA sequences and target genes is available in **Dataset S6**.

### Construction of a genome-wide sgRNA plasmid pool

An oligo pair was designed for each of 3,351 sgRNAs, and 6,702 oligonucleotides were manufactured by IDT (**Dataset S6**), corresponding to two partially anti-complementary 24 nt oligos per annotated ORF at a final concentration of 1 pmol/oligo (oPools™). The annealing process of the oligo pairs in pool resulted in the formation of a double stranded DNA fragment with 20 nt sgRNA base-pairing spacer sequence with a 4 nt 5’ overhang, as described above. Next, the annealing product was 5’ phosphorylated with T4 polynucleotide kinase (New England BioLabs, M0201S) for 40 min at 37°C, and inactivated by incubating at 65°C for 20 min. The two 4 nt overhangs were designed to align with the two adhesive ends of the *BsmB*I-digested pPEPZHi-*mCherry*. To guarantee *mCherry* elimination, a digestion was performed with the *Rrs*II restriction enzyme, which has a cutting site within the *mCherry* gene. The *Rrs*II digestion product was used as a template for PCR with primers OVL6185 and OVL6186, and purified using NucleoSpin Gel and PCR Clean-up Kit. The purified PCR product was digested with *BsmB*I and ligated to the phosphorylated pool-annealing product in a one step process performed in a thermocycler, for 1.5 min at 37°C, 66 cycles of 3 min at 16°C and 5 min at 37°C, followed by 10 min at 80°C, to generate a pPEPZHi-sgRNAlibrary plasmid pool. The pPEPZHi-sgRNAlibrary plasmid pool was transformed into electrocompetent *E. coli* Stbl3 cells. Spec^R^ transformants were pooled, collected and stored at -80°C in LB-glycerol 20%. Collected transformant pools were used for plasmid pool mini-prep purification.

### Construction of CRISPRi library in *H. influenzae*

The purified pPEPZHi-sgRNAlibrary plasmid pool was used to generate a genome-wide CRISPRi library in the *dcas9* strain. The purified sgRNA plasmid pool was separately digested with (i) *Hae*II (New England BioLabs, R0107S) and *Rsr*II (New England BioLabs, R0501S); (ii) *Mme*I (New England BioLabs, R0637S) and *Rsr*II, for 1 h at 37°C, and restriction enzymes were further inactivated for 20 min at 80°C or 65°C, respectively. This generates a pool of linear DNA fragments, mixed together in equal proportion (∼60 µL of the digestion product), and used for transformation of naturally competent *dcas9* bacterial cells using the MI-V method. After the incubation period, the entire volume was plated on HTM-agar with Spec_50_ (300 µL/plate on 150 mm petri dishes), and plates were incubated at 37°C overnight with 5% CO_2_. Pooled transformants were collected with 5 mL sBHI and stored at -80°C in sBHI-glycerol 20%.

### CRISPRi-seq essentiality screen

The *dcas9* CRISPRi library was grown in 25 mL fresh sBHI or CDM medium using 500 μL aliquots of the previously frozen CRISPRi library. Cultures were grown at 37°C with shaking (180 r.p.m) to OD_600_=0.3, and stored (2.5 mL culture + 750 μL 80% glycerol) at -80°C, named working stocks. Library growth for CRISPRi-seq assay was performed using the working stocks. When appropriate, 50 ng/mL aTc was added for induction. All conditions were performed in quadruplicate. For the 7 generations experiments, 250 µL of working stock were inoculated in 24.75 mL medium in the absence (control group) or presence (experimental group) of aTc 50 ng/mL. Cultures were incubated at 37°C with shaking (180 r.p.m) to OD_600_=0.3. Ten mL/culture were next centrifuged (4,000 g, 10 min, 4°C), and pellets were kept at -80°C for subsequent genomic DNA extraction. For 14 generations experiments, the previous cultures were 1:100 diluted (250 µL culture in 24,75 mL medium), and incubated at 37°C, 180 r.p.m. When OD_600_ reached 0.3, 10 mL/culture were harvested by centrifugation (4,000 g, 10 min, 4°C), and pellets were kept as above. For 21 generations assays, cultures from the 14 generations assay were 1:100 diluted as before, incubated at 37°C and 180 r.p.m to reach OD_600_=0.3, 10 mL/culture were centrifuged (4,000 g, 10 min, 4°C), and pellets were kept at -80°C.

### Library preparation and Illumina sequencing

Samples were used for genomic DNA extraction and purification with DNeasy Blood & Tissue kit (Qiagen®, 69506), resuspended in 200 µL DNase-free water/sample, and quantified with a Nanodrop spectrophotometer. Genomic DNA samples served as a template for a one step PCR reaction with primers including barcode index (N5 and N7 illumina series). PCR products were purified from gel extraction, quantified (Qubit® dsDNA HS Assay Kit; life Technologies, Q32851) and assembled in equimolarity for library pooling deep sequencing. Purified amplicons were sequenced on an Aviti systems (Element Biosciences) using single ends 150 bp reads protocols and read primer 1. Data available as a BioProject, ID: PRJNA1274682.

### CRISPRi-seq differential enrichment analysis

Sequencing results were demultiplexed and processed through 2FAST2Q (available at https://github.com/veeninglab/2FAST2Q) to elicit sgRNA counts per sample. Differential enrichment analysis using the R package DESeq2 was performed for evaluation of fitness cost of each sgRNA, as previously described. EggNOG mapper v2 (http://eggnog-mapper.embl.de/) (42) was used to allocate COG categories (one letter code) to each genetic feature of the prokka annotated genome of RdKW20. Enrichment pathway analyses were performed with the R packages ClusterProfiler and enrichlot. Metabolic maps were generated with the R package pathview.

### Generation of Δ*serA* mutant strain

To inactivate the *serA* gene in the RdKW20 strain, a 1,240 bp DNA fragment corresponding to the *serA* coding sequence was PCR amplified (*Phusion* DNA Polymerase) using RdKW20 genomic DNA as template with primers f1_serA_new/2333 and r1_serA_new/2334. This amplicon was cloned into pJET1.2/blunt, generating pJET1.2-*serA/*P1259, which was then linearized by inverse PCR using primers serA_F2_NEW/2376 and serA_R2_NEW/2377 to disrupt the *serA* sequence, followed by ligation to a blunt-ended Spec^R^ gene obtained from pRSM2832 by *EcoR*V digestion, to generate pJET1.2-*serA*::*spec* (P1260). The *serA::spec* disruption cassette (2,410 bp) was PCR amplified with primers f1_serA_new/2333 and r1_serA_new/2334, and used for RdKW20 natural transformation using the MI-V method (38). The Δ*serA::spec* (P1261) strain was selected on sHTM agar with Spec_50_ and confirmed by PCR.

### Microscopy imaging and image processing

Cells were grown for 12 h in 50 mL-falcon tubes with 10 mL sBHI at 37°C with shaking (120 r.p.m). They were next diluted in fresh sBHI to OD_600_=0.05 and incubated at 37°C with shaking (120 r.p.m) for 3 h, in the presence or absence of aTc. When stated, cells were stained with DAPI (Sigma Aldrich, D9542). Briefly, 500 µL of cells were incubated at 37°C with DAPI 100 µg/mL for 15 min, and next centrifuged (5,000 r.p.m. for 5 min) and washed with 1 mL sBHI. Cells were imaged by adding 1 μL of cell suspension onto 1% (w/v) PBS-agarose gel pads. Microscopy was conducted using a Leica DMi8 microscope with a sCMOS DFC9000 GTC (Leica) camera and a ×100/1.40 oil-immersion objective (0.09 working distance). Phase-contrast images were acquired using transmission light with a 100 ms exposure. Snapshot fluorescence images were acquired with a 100 ms exposure, a 405 nm excitation laser module and a DAPI 390 filter (Ex: 395/25 nm, BS: LP 425 Leica 11533333, Em: 460/50 nm). All microscope images were acquired with LasX v.3.4.2.18368 (Leica), and processed with FIJI v2.14.0 to adjust contrast and brightness (43).

### HaemoBrowse

The HaemoBrowse genome browser for *H. influenzae* was created using JBrowse 2 (44), through methods described before (45). In short, the genomes of 86-028NP, NTHi375, R2866, RdKW20 (Refseq accession numbers GCF_000012185.1, GCF_000767075.1, GCF_000165525.1, and GCF_000027305.1, respectively) were acquired from the NCBI, and annotated *de novo* using Prokka (40). The protein sequences were used to generate links to UniProt (46), AlphaFold (47), FoldSeek (48) and STRING (49) for each protein coding sequence, by matching through BLAST (50). Translated protein sequences from strain RdKW20 were also used to match the Prokka annotations to the reference locus tags for RdKW20 (HI_XXXX). Prokka annotation information was used for links to external databases on enzyme classifiers (Enzyme Commission (EC) numbers, and Clusters of Orthologous Groups (COGs). The designed sgRNAs targeting each genome are available from each genome’s sgRNA track. Conservation was determined using panaroo (51) from the Prokka-produced gff-files, through standard settings and “clean-mode” set to “strict”. Genome alignment was determined using MashMap3 (v3.1.3) (52). The NucContent plugin for JBrowse2 is used to provide a track to calculate and visualize GC content (available from https://github.com/jjrozewicki/jbrowse2-plugin-nuccontent).

## Acknowledgments

We thank Drs. Begoña Euba, Nahikari López-López and Javier Asensio-López for helping to define mCDM and L-serine specificities needed for this work.

## Funding

C.G.-C. was funded by a PhD studentship from AEI, PRE2019-088382 and by a FEMS mobility grant. J.M. received funding from the European Union’s Horizon 2020 research and innovation program (Marie Skłodowska-Curie grant N°101018461). A.D.-S.P. is funded by a PhD studentship from AEI, PRE2022-102925. A.B.J. was supported through a Postdoctoral Fellowship grant (TMPFP3_210202) from the Swiss National Science Foundation (SNSF), and a Robert Austrian Research Award from the International Society of Pneumonia and Pneumococcal Diseases. Work at J.W.V. laboratory was supported by the SNSF grants 310030_192517, 310030_200792 and NCCR 51NF40_180541. Work at J.G. laboratory was supported by grants from the Spanish AEI (PID2021-125947OB-I00), SEPAR (875/2019), Regional Govern o Navarra (PC150 and PC136), and from CIBER - an initiative from Instituto de Salud Carlos III (ISCIII), Madrid, Spain.

## References

1. Peltola H. 2000. Worldwide *Haemophilus influenzae* type b disease at the beginning of the 21st century: global analysis of the disease burden 25 years after the use of the polysaccharide vaccine and a decade after the advent of conjugates. Clin Microbiol Rev 13:302–317.

2. Ahearn CP, Gallo MC, Murphy TF. 2017. Insights on persistent airway infection by non-typeable *Haemophilus influenzae* in chronic obstructive pulmonary disease. Pathog Dis 75:1–18.

3. Duell BL, Su YC, Riesbeck K. 2016. Host–pathogen interactions of nontypeable *Haemophilus influenzae*: from commensal to pathogen. FEBS Lett 590:3840–3853.

4. Jalalvand F, Riesbeck K. 2018. Update on non-typeable *Haemophilus influenzae*-mediated disease and vaccine development. Expert Rev Vaccines 17:503–512.

5. Cain AK, Barquist L, Goodman AL, Paulsen IT, Parkhill J, van Opijnen T. 2020. A decade of advances in transposon-insertion sequencing. Nat Rev Genet 21:526–540.

6. Langereis JD, de Jonge MI, Weiser JN. 2014. Binding of human factor H to outer membrane protein P5 of non-typeable *Haemophilus influenzae* contributes to complement resistance. Mol Microbiol 94:89–106.

7. Langereis JD, Weiser JN. 2014. Shielding of a lipooligosaccharide IgM epitope allows evasion of neutrophil-mediated killing of an invasive strain of nontypeable *Haemophilus influenzae*. MBio 5:1–10.

8. Langereis JD, Zomer A, Stunnenberg HG, Burghout P, Hermans PWM. 2013. Nontypeable *Haemophilus influenzae* carbonic anhydrase is important for environmental and intracellular survival. J Bacteriol 195:2737–2746.

9. Larson MH, Gilbert LA, Wang X, Lim WA, Weissman JS, Qi LS. 2013. CRISPR interference (CRISPRi) for sequence-specific control of gene expression. Nat Protoc 8:2180–2196.

10. Bikard D, Jiang W, Samai P, Hochschild A, Zhang F, Marrafini LA. 2013. Programmable repression and activation of bacterial gene expression using and engineered CRISPR-Cas system. Nuclei Acids Res 41(15):7429–37.

11. Qi LS, Larson MH, Gilbert LA, Doudna JA, Weissman JS, Arkin AP, Lim WA. 2013. Repurposing CRISPR as an RNA-guided platform for sequence-specific control of gene expression. Cell 152:1173–1183.

12. Gager C, Flores-mireles AL, Gager C, Flores-mireles AL. 2024. Blunted blades: new CRISPR-derived technologies to dissect microbial multi-drug resistance and biofilm formation. mSphere 9(4):e0064223.

13. Ghavami S, Pandi A. 2021. CRISPR interference and its applications. Progress in Molecular Biology and Translational Science, 1st ed. Elsevier Inc.

14. Peters JM, Colavin A, Shi H, Czarny TL, Larson MH, Wong S, Hawkins JS, Lu CHS, Koo BM, Marta E, Shiver AL, Whitehead EH, Weissman JS, Brown ED, Qi LS, Huang KC, Gross CA. 2016. A comprehensive, CRISPR-based functional analysis of essential genes in bacteria. Cell 165:1493–1506.

15. de Bakker V, Liu X, Bravo AM, Veening J-W. 2022. CRISPRi-seq for genome-wide fitness quantification in bacteria. Nat Protoc 17:252–281.

16. Liu X, Gallay C, Kjos M, Domenech A, Slager J, van Kessel SP, Knoops K, Sorg RA, Zhang J-R, Veening J-W. 2017. High-throughput CRISPRi phenotyping identifies new essential genes in *Streptococcus pneumoniae*. Mol Syst Biol 13:931.

17. Liu X, Bakker V De, Heggenhougen MV, Mårli MT, Frøynes AH, Salehian Z, Porcellato D, Angeles DM, Veening J, Kjos M. 2024. Genome-wide CRISPRi screens for high-throughput fitness quantification and identification of determinants for dalbavancin susceptibility in *Staphylococcus aureus*. mSystems 9(7):e0128923.

18. Liu X, Kimmey JM, Matarazzo L, de Bakker V, Van Maele L, Sirard J-C, Nizet V, Veening J-W. 2021. Exploration of bacterial bottlenecks and *Streptococcus pneumoniae* pathogenesis by CRISPRi-Seq. Cell Host Microbe 29:107–120.e6.

19. de Wet TJ, Winkler KR, Mhlanga M, Mizrahi V, Warner DF. 2020. Arrayed CRISPRi and quantitative imaging describe the morphotypic landscape of essential mycobacterial genes. Elife 9:1–36.

20. Wang T, Guan C, Guo J, Liu B, Wu Y, Xie Z, Zhang C, Xing XH. 2018. Pooled CRISPR interference screening enables genome-scale functional genomics study in bacteria with superior performance-net. Nat Commun 9:2475.

21. Rousset F, Cui L, Siouve E, Becavin C, Depardieu F, Bikard D. 2018. Genome-wide CRISPR-dCas9 screens in *E. coli* identify essential genes and phage host factors. PLoS Genet 14:1–28.

22. Lee HH, Ostrov N, Wong BG, Gold MA, Khalil AS, Church GM. 2019. Functional genomics of the rapidly replicating bacterium *Vibrio natriegens* by CRISPRi. Nat Microbiol 4:1105–1113.

23. Minhas V, Domenech A, Synefiaridou D, Straume D, Brendel M, Cebrero G, Liu X, Costa C, Baldry M, Sirard JC, Perez C, Gisch N, Hammerschmidt S, Håvarstein LS, Veening JW. 2023. Competence remodels the pneumococcal cell wall exposing key surface virulence factors that mediate increased host adherence. PLoS Biol. 21(1):e3001990.

24. Liu X, Van Maele L, Matarazzo L, Soulard D, Alves Duarte da Silva V, de Bakker V, Dénéréaz J, Bock FP, Taschner M, Ou J, Gruber S, Nizet V, Sirard JC, Veening JW. 2014. A conserved antigen induces respiratory Th17-mediated broac serotype protection against pneumococcal superinfection. Cell Host Microbe. 2024. 32(3):304–314.e8.

25. Mårli MT, Nordraak AOO, de Bakker V, Winther AR, Liu X, Veening JW, Porcellato D, Kjos M. 2025. Genome-wide analysis of fitness determinants of *Staphylococcus aureus* during growth in milk. PLoS Pathog. 21(4):e1013080.

26. Zhang Y, Zhang T, Xiao X, Wang Y, Kawalek A, Ou J, Ren A, Sun W, de Bakker V, Liu Y, Li Y, Yang L, Ye L, Jia N, Veening JW, Liu X. 2025. CRISPRi screen identifies FprB as a synergistic target for gallium therapy in *Pseudomonas aeruginosa*. Nat Commun. 16(1):5870.

27. Sorg RA, Gallay C, Van Maele L, Sirard J-C, Veening J-W. 2020. Synthetic gene-regulatory networks in the opportunistic human pathogen *Streptococcus pneumoniae*. Proc Natl Acad Sci U S A 117:27608–27619.

28. Wong SM, Akerley BJ. 2003. Inducible expression system and marker-linked mutagenesis approach for functional genomics of *Haemophilus influenzae*. Gene 316:177–86.

29. Morey P, Viadas C, Euba B, Hood DW, Barberán M, Gil C, Grilló MJ, Bengoechea JA, Garmendia J. 2013. Relative contributions of lipooligosaccharide inner and outer core modifications to nontypeable *Haemophilus influenzae* pathogenesis. Infect Immun. 81(11):4100–11.

30. López-López N, León DS, de Castro S, Díez-Martínez R, Iglesias-Bexiga M, Camarasa MJ, Menéndez M, Nogales J, Garmendia J. 2022. Interrogation of essentiality in the reconstructed *Haemophilus influenzae* metabolic network identifies lipid metabolism antimicrobial targets: preclinical evaluation of a FabH β-Ketoacyl-ACP synthase inhibitor. mSystems 7:e0145921.

31. Gawronski JD, Wong SMS, Giannoukos G, Ward D V., Akerley BJ. 2009. Tracking insertion mutants within libraries by deep sequencing and a genome-wide screen for *Haemophilus* genes required in the lung. Proc Natl Acad Sci U S A 106:16422–16427.

32. Wong SM, Bernui M, Shen H, Akerley BJ. 2013. Genome-wide fitness profiling reveals adaptations required by *Haemophilus* in coinfection with influenza A virus in the murine lung. Proc Natl Acad Sci U S A 110:15413–15418.

33. Othman DSMP, Schirra H, McEwan AG, Kappler U. 2014. Metabolic versatility in *Haemophilus influenzae*: A metabolomic and genomic analysis. Front Microbiol 5:1– 10.

34. López-López N, Euba B, Hill J, Dhouib R, Caballero L, Leiva J, Hosmer J, Cuesta S, Ramos-Vivas J, Díez-Martínez R, Schirra HJ, Blank LM, Kappler U, Garmendia J. 2020. *Haemophilus influenzae* glucose catabolism leading to production of the immunometabolite acetate has a key contribution to the host airway-pathogen interplay. ACS Infect Dis 6:406–421.

35. Nogales J, Garmendia J. 2022. Bacterial metabolism and pathogenesis intimate intertwining: time for metabolic modelling to come into action. Microb Biotechnol 15:95–102.

36. Mobegi FM, Van Hijum SA, Burghout P, Bootsma HJ, De Vries SP, Van Der Gaast-de Jongh CE, Simonetti E, Langereis JD, Hermans PW, De Jonge MI, Zomer A. 2014. From microbial gene essentiality to novel antimicrobial drug targets. BMC Genomics 15:958.

37. Gawronski JD, Wong SM, Giannoukos G, Ward DV, Akerley BJ. 2009. Tracking insertion mutants within libraries by deep sequencing and a genome-wide screen for *Haemophilus influenzae* genes required in the lung. Proc Natl Acad Sci USA. 106(38):16422–7.

38. Herriott RM, Meyer EM, Vogt M. 1970. Defined nongrowth media for stage II development of competence in *Haemophilus influenzae*. J Bacteriol 101:517–524.

39. Fernández-Calvet A, Rodríguez-Arce I, Almagro G, Moleres J, Euba B, Caballero L, Martí S, Ramos-Vivas J, Bartholomew TL, Morales X, Ortíz-De-Solórzano C, Yuste JE, Bengoechea JA, Conde-Álvarez R, Garmendia J. 2018. *Modulation of* Haemophilus influenzae interaction with hydrophobic molecules by the VacJ/MlaA lipoprotein impacts strongly on its interplay with the airways. Sci Rep 8:1–17.

40. Seemann T. 2014. Prokka: rapid prokaryotic genome annotation. Bioinformatics 30:2068–2069.

41. Bjånes E, Stream A, Janssen AB, Gibson PS, Bravo AM, Dahesh S, Baker JL. 2024. An efficient *in vivo*-inducible CRISPR interference system for group A *Streptococcus* genetic analysis and pathogenesis studies. mBio 15(8):e0084024.

42. Cantalapiedra CP, Hernández-Plaza A, Letunic I, Bork P, Huerta-Cepas J. 2021. eggNOG-mapper v2: Functional Annotation, Orthology Assignments, and Domain Prediction at the Metagenomic Scale. Mol Biol Evol. United States.

43. Schindelin J, Arganda-Carreras I, Frise E, Kaynig V, Longair M, Pietzsch T, Preibisch S, Rueden C, Saalfeld S, Schmid B, Tinevez JY, White DJ, Hartenstein V, Eliceiri K, Tomancak P, Cardona A. 2012. Fiji: an open-source platform for biological-image analysis. Nat Methods 9:676–682.

44. Diesh C, Stevens GJ, Xie P, De Jesus Martinez T, Hershberg EA, Leung A, Guo E, Dider S, Zhang J, Bridge C, Hogue G, Duncan A, Morgan M, Flores T, Bimber BN, Haw R, Cain S, Buels RM, Stein LD, Holmes IH. 2023. JBrowse 2: a modular genome browser with views of synteny and structural variation. Genome Biol 24.

45. Janssen AB, Gibson PS, Bravo AM, De Bakker V, Slager J, Veening J-W. 2024. PneumoBrowse 2: An integrated visual platform for curated genome annotation and multiomics data analysis of *Streptococcus pneumoniae*. Nucleic Acids Res 53(D1):D839–D851.

46. Bateman A, Martin M-J, Orchard S, Magrane M, Agivetova R, Ahmad S, Alpi E, Bowler-Barnett EH, Britto R, Bursteinas B, Bye-A-Jee H, Coetzee R, Cukura A, Da Silva A, Denny P, Dogan T, Ebenezer T, Fan J, Castro LG, Garmiri P, Georghiou G, Gonzales L, Hatton-Ellis E, Hussein A, Ignatchenko A, Insana G, Ishtiaq R, Jokinen P, Joshi V, Jyothi D, Lock A, Lopez R, Luciani A, Luo J, Lussi Y, MacDougall A, Madeira F, Mahmoudy M, Menchi M, Mishra A, Moulang K, Nightingale A, Oliveira CS, Pundir S, Qi G, Raj S, Rice D, Lopez MR, Saidi R, Sampson J, Sawford T, Speretta E, Turner E, Tyagi N, Vasudev P, Volynkin V, Warner K, Watkins X, Zaru R, Zellner H, Bridge A, Poux S, Redaschi N, Aimo L, Argoud-Puy G, Auchincloss A, Axelsen K, Bansal P, Baratin D, Blatter M-C, Bolleman J, Boutet E, Breuza L, Casals-Casas C, De Castro E, Echioukh KC, Coudert E, Cuche B, Doche M, Dornevil D, Estreicher A, Famiglietti ML, Feuermann M, Gasteiger E, Gehant S, Gerritsen V, Gos A, Gruaz-Gumowski N, Hinz U, Hulo C, Hyka-Nouspikel N, Jungo F, Keller G, Kerhornou A, Lara V, Le Mercier P, Lieberherr D, Lombardot T, Martin X, Masson P, Morgat A, Neto TB, Paesano S, Pedruzzi I, Pilbout S, Pourcel L, Pozzato M, Pruess M, Rivoire C, Sigrist C, Sonesson K, Stutz A, Sundaram S, Tognolli M, Verbregue L, Wu CH, Arighi CN, Arminski L, Chen C, Chen Y, Garavelli JS, Huang H, Laiho K, McGarvey P, Natale DA, Ross K, Vinayaka CR, Wang Q, Wang Y, Yeh L-S, Zhang J, Ruch P, Teodoro D. 2021. UniProt: the universal protein knowledgebase in 2021. Nucleic Acids Res 49:D480–D489.

47. Jumper J, Evans R, Pritzel A, Green T, Figurnov M, Ronneberger O, Tunyasuvunakool K, Bates R, Žídek A, Potapenko A, Bridgland A, Meyer C, Kohl SAA, Ballard AJ, Cowie A, Romera-Paredes B, Nikolov S, Jain R, Adler J, Back T, Petersen S, Reiman D, Clancy E, Zielinski M, Steinegger M, Pacholska M, Berghammer T, Bodenstein S, Silver D, Vinyals O, Senior AW, Kavukcuoglu K, Kohli P, Hassabis D. 2021. Highly accurate protein structure prediction with AlphaFold. Nature 596:583–589.

48. Van Kempen M, Kim SS, Tumescheit C, Mirdita M, Lee J, Gilchrist CLM, Söding J, Steinegger M. 2024. Fast and accurate protein structure search with Foldseek. Nat Biotechnol 42:243–246.

49. Szklarczyk D, Kirsch R, Koutrouli M, Nastou K, Mehryary F, Hachilif R, Gable AL, Fang T, Doncheva NT, Pyysalo S, Bork P, Jensen LJ, von Mering C. 2023. The STRING database in 2023: protein–protein association networks and functional enrichment analyses for any sequenced genome of interest. Nucleic Acids Res 51:D638–D646.

50. AltschuP SF, Gish W, Miller W, Myers EW, Lipman DJ. 1990. Basic Local Alignment Search Tool. J Mol Biol 215:403–410.

51. Tonkin-Hill G, MacAlasdair N, Ruis C, Weimann A, Horesh G, Lees JA, Gladstone RA, Lo S, Beaudoin C, Floto RA, Frost SDW, Corander J, Bentley SD, Parkhill J. 2020. Producing polished prokaryotic pangenomes with the Panaroo pipeline. Genome Biol 21:1–21.

52. Kille B, Garrison E, Treangen TJ, Phillippy AM. 2023. Minmers are a generalization of minimizers that enable unbiased local Jaccard estimation. Bioinformatics 39(9):btad512.

